# Quantitative crosslinking and mass spectrometry determine binding interfaces and affinities mediating kinetochore stabilization

**DOI:** 10.1101/2022.03.31.486303

**Authors:** Goetz Hagemann, Victor Solis-Mezarino, Sylvia Singh, Mia Potocnjak, Chandni Kumar, Franz Herzog

## Abstract

Crosslinking and mass spectrometry (XLMS) are used in integrative structural biology to acquire spatial restraints. We found a dependency between crosslink distances and intensities and developed a quantitative workflow to simultaneously estimate apparent dissociation constants (K_D_) of contacts within multi-subunit complexes and to aid interface prediction. Quantitative XLMS was applied to study the assembly of the macromolecular kinetochore complex, which is built on centromeric chromatin and establishes a stable link to spindle microtubules in order to segregate chromosomes during cell division. Inter-protein crosslink intensities facilitated determination of phosphorylation-induced binding interfaces and affinity changes. Phosphorylation of outer and inner kinetochore proteins mediated cooperative kinetochore stabilization and decreased the K_D_ values of its interactions to the centromeric nucleosome by ~200-fold, which was essential for cell viability. This work demonstrates the potential of quantitative XLMS for characterizing mechanistic effects on protein assemblies upon post-translational modifications or cofactor interaction and for biological modeling.

## Main

Distance restraints derived from the mass spectrometric identification of crosslinked amino acids (XLMS) are widely applied in integrative approaches to determine protein connectivity^1^ and to model the topology of proteins and their domains in a complex^2^. Quantification of crosslinks has been initially implemented to detect conformational changes and domain interactions^3–5^. Besides structure, the critical determinant of the molecular mechanism of a complex is the interaction strength of its subunit contacts, which can be modulated through cofactors or post-translational modifications to execute its biological function on time. Several biophysical methods^6^ are available to measure protein-protein affinity through estimation of the apparent dissociation constant (K_D_), but the individual methods mainly analyze binary interactions and require high protein concentrations, protein engineering, immobilization or labeling which may affect the integrity of complexes. We reasoned that crosslink intensities provide a quantitative measure for the formed complex and the free subunits at the equilibrium state. Thus, we investigated whether crosslink intensities facilitate the simultaneous estimation of individual protein-protein affinities within kinetochore multi-subunit complexes.

The kinetochore is a macromolecular protein complex assembled at centromeric chromatin that ensures the fidelity of chromosome segregation by connecting chromosomes and spindle microtubules and by integrating feedback control mechanisms^7,8^. In order to bi-orient chromosomes on the mitotic spindle the budding yeast kinetochore has to transmit forces of ~10 pN ^9,10^ by forming a load-bearing attachment to spindle microtubules and a high-affinity link to the centromeric nucleosome, marked by the histone H3 variant Cse4^CENP-A^ (human orthologs are superscripted if appropriate). The kinetochore subunits are largely conserved between budding yeast and humans^11,12^ and form stable subcomplexes, which are organized in two layers of the kinetochore architecture. The outer kinetochore, a 10-subunit network that is built up on the inner kinetochore, forms the microtubule binding site. The inner kinetochore is assembled by at least 15 proteins on centromeric chromatin with Mif2 and Ame1/Okp1 directly linking the outer kinetochore MTW1 (Mtw1/Nnf1/Dsn1/Nsl1) complex to the Cse4-NCP (Cse4 containing nucleosome core particle) in budding yeast^13–16^. Whereas the human kinetochore assembly is temporally regulated, establishing a microtubule attachment site in mitosis, budding yeast kinetochores are built up and attached to a single microtubule almost throughout the entire cell cycle^7,17,18^. In both species, phosphorylation of Dsn1^DSN1^ by the mitotic kinase Ipl1^Aurora-B^ stabilizes the recruitment of the outer to the inner kinetochore^19–21^. In addition, phosphorylation of the human kinetochore by Plk1 has been shown to stabilize the inner kinetochore architecture at centromeric chromatin to withstand the pulling forces of depolymerizing microtubules^22^.

By quantifying crosslink-derived restraints we found a dependency between crosslink distances and intensities. This relation was applied to improve the prediction of protein binding interfaces and to determine apparent K_D_ values of their interactions, which provided quantitative measures to capture different functional states of the kinetochore. Our approach facilitated the detection of phosphorylation-induced changes in binding affinities between the centromeric nucleosome and a minimal kinetochore assembly composed of the outer kinetochore MTW1^MIS12^ complex, the inner kinetochore Mif2^CENP-C^ and Ame1/Okp1^CENP-U/Q^ proteins.

## Results

### Determination of crosslink intensity and its dependence on crosslink distance

To quantify protein crosslinks, we first extracted the MS1 peak intensities of the MS2 based crosslink identifications using an in-house bioinformatics pipeline that merges the open-source software tools xQuest/xProphet^23,24^ and OpenMS^25^ (Fig. 1 and Methods). Protein complexes were crosslinked by modifying the α-amino groups with the isotopically labeled BS2G-d_0_/d_6_ reagent and crosslinked peptide fractions were analyzed by liquid chromatography coupled to tandem mass spectrometry. The raw files were processed by the xQuest/xProphet software to identify the crosslinked peptides, their precursor ion masses and retention times. This information was subsequently used for the extraction of ion chromatograms by the OpenMS software tool, which were summarized in text tables. The quantification pipeline was benchmarked against available datasets showing that our bioinformatics workflow performs similarly to previously reported software tools in terms of signal detection rate and accuracy of quantification, and is independent of the crosslinker type (Supplementary Fig. 1).

**Fig. 1:**
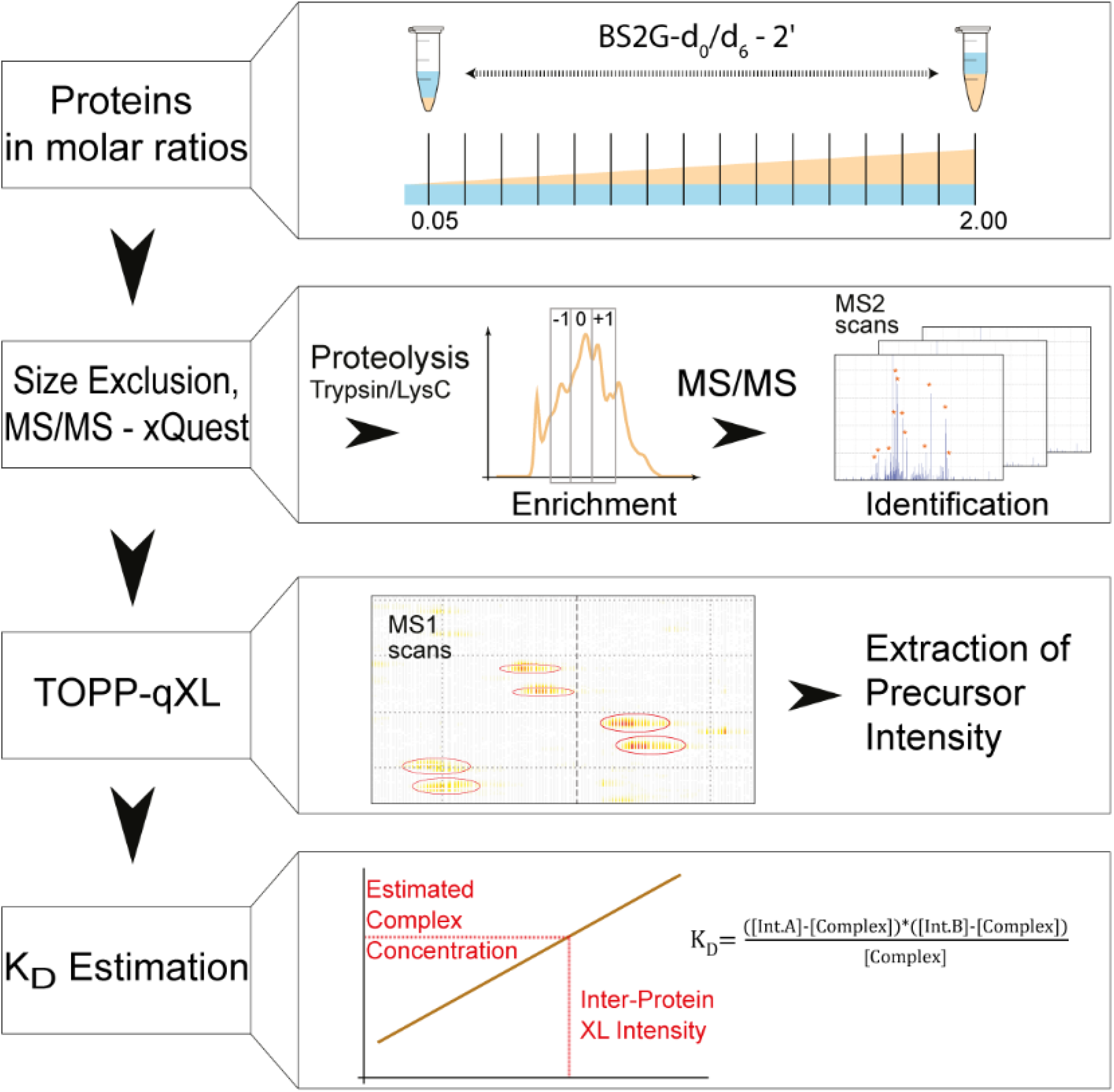
Schematic workflow of estimating protein affinities by quantitative XLMS. The binding partners were titrated by increasing the molar ratio of one interactor. Crosslinked proteins were proteolytically digested, enriched by size exclusion chromatography and linked peptides were identified by tandem mass spectrometry and the software xQuest^23,24^. Precursor intensities of the crosslinks were extracted using our TOPP-qXL (The OpenMS Proteomics Pipeline-quantitative XLMS) bioinformatics workflow. The intensities of intra- and inter-protein site-site links were applied to estimate the concentration of free interactors and complex and for the statistical modeling of apparent K_D_ values.

Quantifying the crosslinks of published multi-protein complex datasets^26,27^ and mapping the corresponding Euclidean lysine-lysine distances on available crystal structures, including those of RNA polymerase I and II, indicated that shorter Euclidean distances between the crosslinked lysines correlate with increasing crosslink intensities (Fig. 2a and Supplementary Fig. 2). We assumed that the inter-protein crosslink intensity is also affected by the physicochemical microenvironment of individual lysines as well as by a competition for the formation of intra-, inter-protein or mono-links at a specific lysine site during the crosslinking reaction. To assess whether crosslink intensities increase for lysine sites proximal to binding interfaces, we mapped the intensity values along the sequences of the RPB1-RPB2 interaction in RNA polymerase II (Supplementary Fig. 3a) as well as of the budding yeast kinetochore Cnn1-Spc24/25 interaction (Supplementary Fig. 3b). We normalized the inter-protein crosslink intensities to the sum of intensities of intra- and inter-protein crosslinks and monolinks occurring at a specific lysine residue. This normalized intensity value or ‘Relative Interface Propensity Index’ (RIPI) served as an indicator for putative interface sequences and was applied in an heuristic approach together with secondary structural elements, sequence conservation and other parameters to aid in the prediction of protein-protein interfaces (Supplementary Fig. 3 and Methods).

**Fig. 2:**
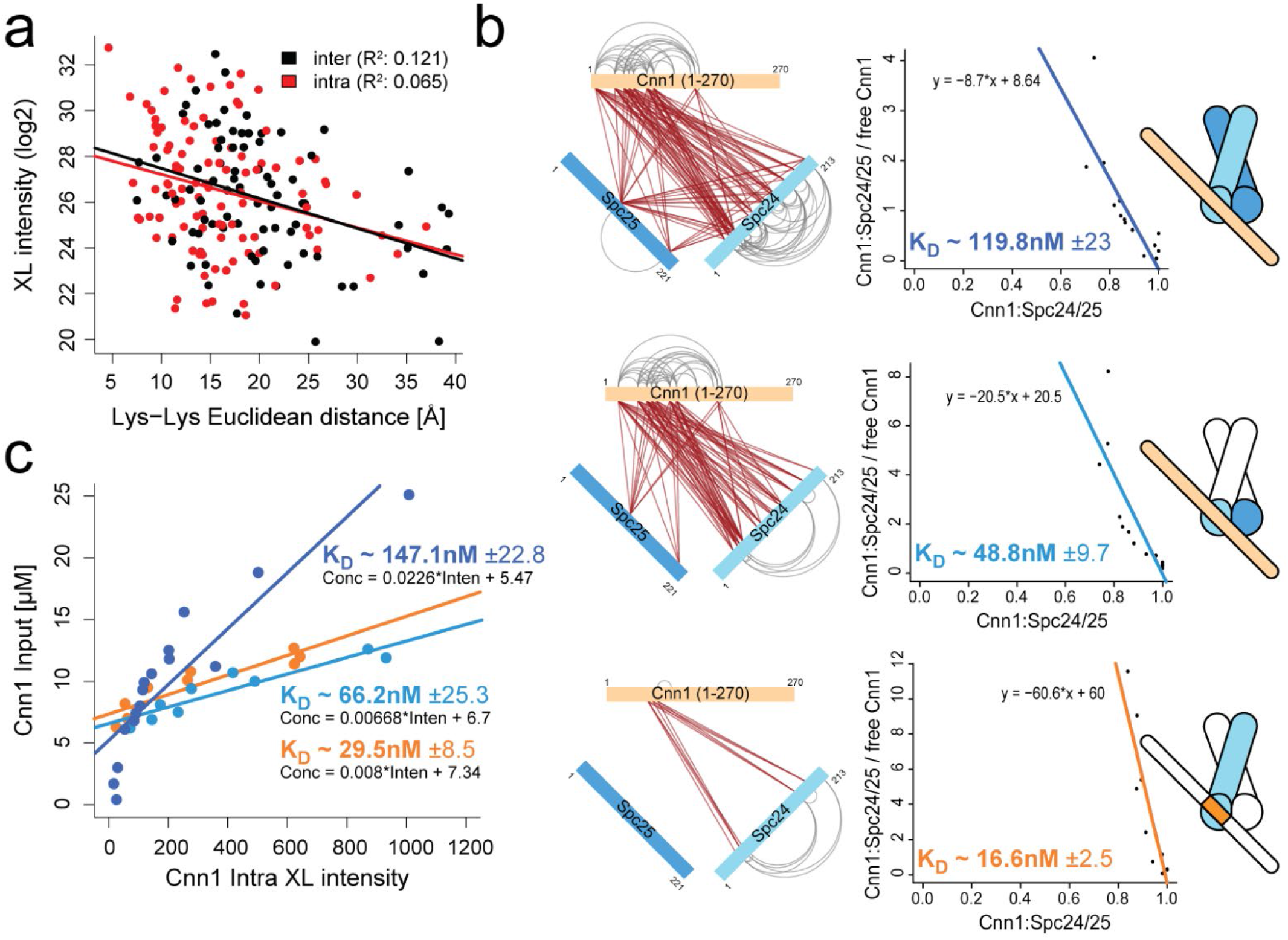
Estimation of apparent K_D_ values in protein complexes using quantitative XLMS. **a**, Correlation of increasing crosslink intensities with decreasing Euclidean distances between crosslinked residues obtained from RNA polymerases analyses (Supplementary Fig. 2). The R-squared statistics and Fisher’s test was computed (p-value(intra)=0.00526, p-value(inter)=0.00098). **b**, Estimation of apparent K_D_ values of the Cnn1^1-270^:Spc24/25 interaction by the *Scatchard* plot using different subsets of inter-protein crosslinks to quantify complex formation. **c**, Apparent K_D_ values were calculated based on the concentration of formed complex interpolated from the linear regression and averaged across molar ratios of the titration steps.

### Estimation of protein affinities based on crosslink intensities

We further applied inter- and intra-protein crosslink intensities to estimate the concentrations of the formed complex and the free subunits according to the steady state equilibrium in solution. To assess whether crosslink intensities supported the estimation of binding affinities we purified recombinant kinetochore subunits and titrated complex formation over a range of molar ratios. First, the inner and outer kinetochore proteins Cnn1^1-270^ and Spc24/25^28^, respectively, were titrated by applying molar ratios from 0.05:1 to 2:1 (Fig. 1, Supplementary Fig. 4 and Supplementary Table 1). To capture the equilibrium state of the binding reaction by crosslinking, the reaction time of the BS2G-d_0_/d_6_ reagent was limited to 2 minutes. Intra-protein crosslink intensities of the constant interactor facilitated the normalization between titration steps and those of the titrated interactor enabled the calculation of a linear regression of the intra-protein intensities on the increasing input protein concentrations (Supplementary Fig. 4 and Supplementary Table 2). The regression model was applied to interpolate the concentration of the formed complex from the inter-protein crosslink intensities (Fig. 1).

The estimation of the apparent K_D_ value was performed first by the *Scatchard* plot^29^ (Fig. 2b and Methods) that indicates the K_D_ value as the negative inverse of the slope. We calculated the K_D_ values for three different sets of inter-protein crosslinks (Fig. 2b). Applying either all inter-protein crosslinks to Cnn1^1-270^ or only those intersecting with the structured domains of Spc24/25 resulted in K_D_ values of 120 nM or 50 nM, respectively. The subset of inter-links decorating the Cnn1^60-84^ motif, that is required for mediating the interaction with Spc24/25, showed a K_D_ of 15 nM which agrees with the value previously obtained by isothermal titration calorimetry (ITC)^28^. This observation is consistent with the notion that residues proximal to the interface may be stably positioned and thus yield relatively higher inter-protein crosslink intensities. The second method used the steady state equilibrium equation to calculate the mean of K_D_ values of each titration step from the concentrations of the formed complex and the free interactors (Figs. 1, 2c and Supplementary Table 3). The second approach based on the steady state equilibrium equation closely reproduced the values obtained by the *Scatchard* plot. Moreover, a similar experiment was performed by titrating increasing concentrations of the Cnn1^60-84^ peptide, containing the minimal binding motif, against the Spc24/25 dimer. The estimated K_D_ value of 2.6 µM (Supplementary Figs. 5 and 6) agrees with previous ITC measurements^28^ and suggests that Cnn1 sequences outside the Cnn1^60-84^ motif contribute to the stabilization of the interaction.

### Phosphorylation of the inner kinetochore by Cdc5^Plk1^ induces its cooperative stabilisation on Cse4 nucleosomes

To determine the apparent K_D_ values of the individual interactions that assemble the kinetochore on the octameric Cse4 nucleosome, we *in vitro* reconstituted kinetochore complexes of up to 11 recombinant proteins (Fig. 3a) purified from *E. coli*, except Mif2, which was isolated from insect cells (Methods). We first reproduced the interaction of Mif2 and Ame1/Okp1^15^, both of which directly bind Cse4-NCPs^13,14,30^, and found that this interaction was lost upon dephosphorylation of Mif2 (Fig. 3b). *In vitro* phosphorylation of lambda-phosphatase-treated Mif2 by the mitotic kinases Cdc28^CDK1^, Cdc5^PLK1^, Ipl1^Aurora-B^ and Mps1^MPS1^ showed that Cdc5^PLK1^ restored Ame1/Okp1 binding to levels detected at insect cell-phosphorylated Mif2 (Fig. 3b). For the subsequent XLMS and binding experiments Mif2 wild-type and mutant proteins were *in vitro* phosphorylated by Cdc5 and are indicated as Mif2^*^.

**Fig. 3:**
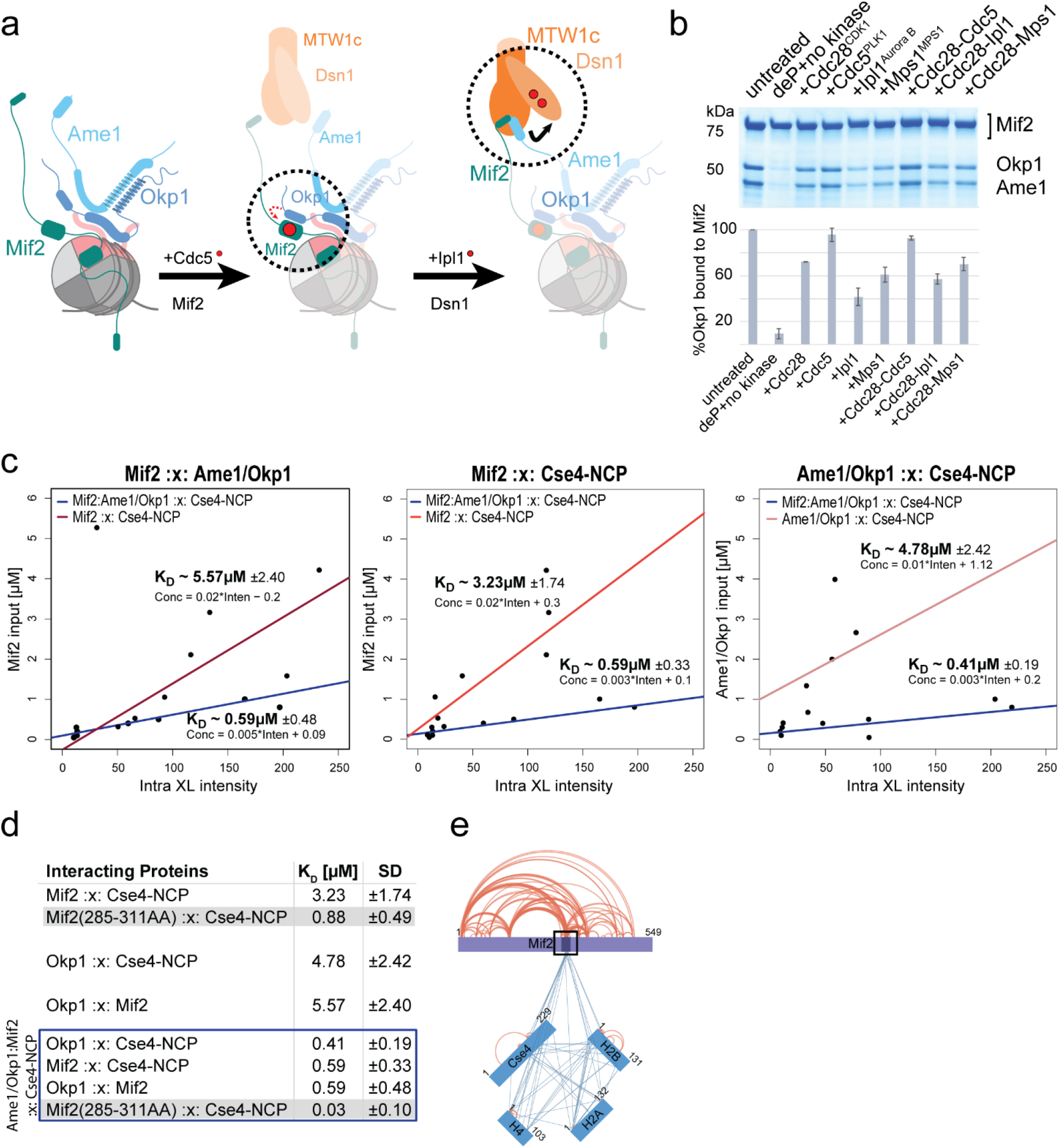
The phosphorylation-dependent binding of Mif2* to Ame1/Okp1 cooperatively stabilizes their interactions with the Cse4-NCP. **a**, Reconstitution of the Mif2:Ame1/Okp1 interaction by dephosphorylation (deP) of Mif2 and subsequent *in vitro* phosphorylation with the indicated kinases (mean ±SD of 3 replicates). **b**, Schematic representation of the assembly of MTW1c, Mif2, and Ame1/Okp1 on the Cse4-NCP. **c**, Estimation of apparent K_D_ values from XLMS analysis of Mif2*:Ame1/Okp1, Mif2*:Cse4-NCP and Ame1/Okp1:Cse4-NCP complexes compared to the apparent K_D_ values within the Mif2*:Ame1/Okp1:Cse4-NCP complex (mean ±SD of 3 replicates). **d**, Summary of estimated K_D_ values including the K_D_ determination of the Mif2:Cse4-NCP interaction using the subset of inter-protein crosslinks to the Mif2^285-311^ signature motif. **e**, Network plot of Mif2*:Cse4-NCP crosslinks intersecting with Mif2^285-311^.

We first estimated apparent K_D_ values of the individual interactions of Cse4-NCP, Mif2^*^ and Ame1/Okp1 by titrating the Cse4-NCP with increasing concentrations of Mif2^*^ or Ame1/Okp1 and by titrating Ame1/Okp1 with Mif2^*^ (Figs. 3c, d, e and Supplementary Fig. 7). The binding affinities of these binary interactions were then compared to the K_D_ values of these interactions in the Mif2^*^:Ame1/Okp1:Cse4-NCP complex. Only intra-and inter-protein crosslinks yielding the extraction of intensities from all 3 replicates (Supplementary Fig. 8) were applied to estimate the apparent K_D_ values based on the steady state equilibrium equation (Supplementary Table 4). The affinities of the binary interactions ranging from 3 to 6 µM were increased 6-fold for the Mif2^*^:Cse4-NCP interaction and 10-fold for the Ame1/Okp1:Cse4-NCP and Mif2^*^:Ame1/Okp1 interactions in the Mif2^*^:Ame1/Okp1:Cse4-NCP complex, indicating cooperative stabilization upon the phosphorylation-induced Mif2^*^:Ame1/Okp1 interaction (Figs. 3c, d and Supplementary Table S5).

Similar to the K_D_ calculation of the Cnn1^1-270^:Spc24/25 interaction, the restriction of inter-protein crosslinks to the subset intersecting with the minimal binding motif, the Mif2^285-311^ signature motif (Figs. 3d and e) which directly binds the CENP-A C-terminus^16,31^, resulted in lower K_D_ values. The K_D_ value of the Mif2^*^:Cse4-NCP complex was reduced from 3.2 to 0.9 µM which is in agreement with ITC measurements of the Mif2^285-311^ peptide with the Cse4-NCP showing a K_D_ of 0.5 µM^31^. Upon the cooperative interactions of Mif2^*^ and Ame1/Okp1 to the Cse4-NCP the K_D_ dropped by a factor of ~30 from 0.6 to 0.03 µM (Figs. 3d and e) demonstrating that quantitative XLMS facilitates the estimation of apparent K_D_ values and the detection of ~200-fold affinity changes in multi-subunit complexes.

### Phosphorylation of outer and inner kinetochore proteins synergistically enhance kinetochore stabilization at the Cse4 nucleosome

The tetrameric MTW1^MIS12^ complex binds Mif2^CENP-C^ and Ame1/Okp1. This interaction is stabilized upon Dsn1^DSN1^ phosphorylation by Ipl1^Aurora-B^ which releases the masking of the Mif2^CENP-C^ and Ame1/Okp1 binding sites at the MTW1^MIS12^ head I domain by Dsn1^DSN1^ (Fig. 3a)^19,20,32,33^. To test whether addition of MTW1c affected the interactions of Cse4-NCP with Mif2^*^ and Ame1/Okp1, we titrated constant levels of Cse4-NCPs with increasing concentrations of an equimolar mixture of Mif2^*^:Ame1/Okp1:MTW1c which contained either wild-type Dsn1 or the phosphorylation-mimicking Dsn1^S240D,S250D^ mutant (Supplementary Fig. 9). The quantification of inter-protein crosslinks (Supplementary Fig. 10 and Supplementary Table 6) intersecting with Mif2 indicated the previously reported Mif2 interfaces to the Cse4-NCP^15,16,31^ and to the MTW1c (Supplementary Figs. 11 and 12a)^8^. The estimation of binding affinities by the steady state equilibrium equation revealed that addition of wild-type MTW1c did not affect the K_D_ values of Mif2^*^ and Ame1/Okp1 to the Cse4-NCP (Figs. 3d, 4a and b and Supplementary Table 7). In comparison, the phosphorylation-mimicking MTW1c(Dsn1^S240D,S250D^) decreased the K_D_ values by ~20-fold and a similar change in affinity was observed for the Mif2:Okp1 interaction (Figs. 4a and b). This indicated that in addition to the Mif2^*^:Okp1 interaction, putatively mediated by Cdc5, phosphorylation of Dsn1 by Ipl1 synergistically enhanced the binding affinity of Mif2^*^ and Ame1/Okp1 to the Cse4-NCP.

**Fig. 4:**
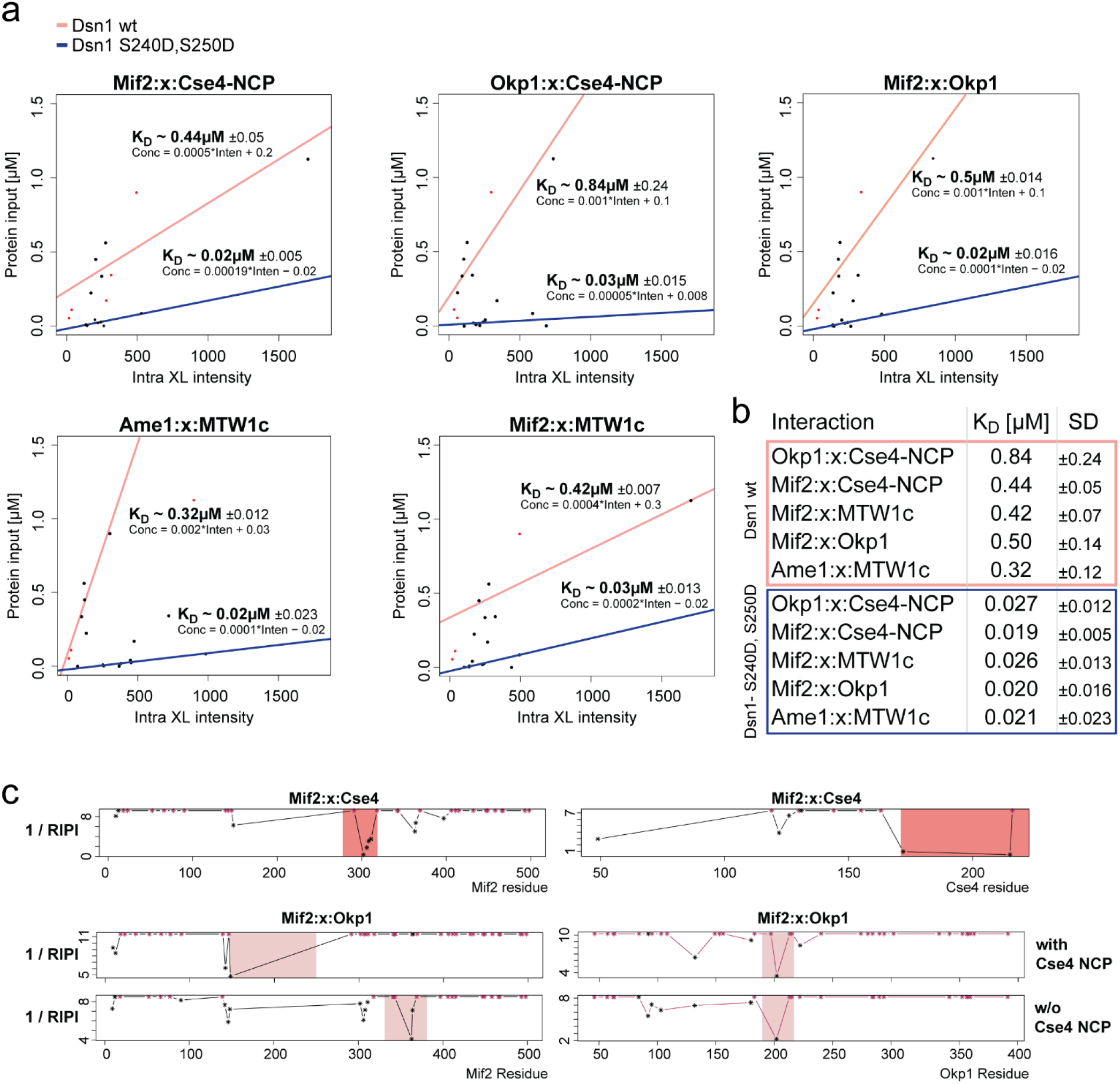
Binding of the MTW1c cooperatively increased the affinity of the Mif2* and Ame1/Okp1 interaction to the Cse4-NCP. **a**, Estimation of apparent K_D_ values by titrating Cse4-NCPs with increasing concentrations of a MTW1c:Mif2*:Ame1/Okp1 complex containing either wild-type Dsn1 or phosphorylation-mimicking Dsn1^S240D,S250D^ (mean ±SD of 2 replicates). **b**, Summary of K_D_ values showing the effect upon binding of MTW1c(Dsn1^S240D,S250D^). **c**, Prediction of the Mif2*:Cse4 and Mif2*:Okp1 interface by calculating the RIPI based on inter-protein crosslink intensities (Supplementary Fig. 2 and Methods).

### The phosphorylation-induced cooperativity mediating kinetochore stabilization is essential in budding yeast

The RIPI calculated from inter-protein crosslink intensities of the Mif2^*^:Okp1 interaction identified Mif2^150-250^ and Okp1^180-220^ as the putative binding motifs (Figs. 3a, 4c and Supplementary Fig. S12b). Based on the indicated regions, mutant proteins were generated to assess the required Mif2 phosphorylation sites mediating its interaction with Ame1/Okp1 in *in vitro* binding and cell viability assays. The Mif2^Δ221-240^ mutant abrogated the Mif2^*^:Ame1/Okp1 interaction *in vitro* whereas Mif2^Δ200-230^ still bound (Fig. 5a). By assessing the phosphorylation dependency of this interaction (Fig. 3b), we found that Ame1/Okp1 binding was lost upon mutating 9 serines to alanines within Mif2^217-240^ (Fig. 5a and Supplementary Fig. 13a). Ectopic expression of the Mif2 mutants, that were impaired in Ame1/Okp1 binding, did not affect growth of budding yeast cells after nuclear depletion of endogenous Mif2 (Supplementary Figs. 14, 15a and Supplementary Table 8). Similarly, the Dsn1^S240A,S250A,S264A^ mutant, which has been previously shown to affect binding of the outer kinetochore MTW1 complex to the inner kinetochore, was viable (Fig. 5b)^19^. Notably, ectopic expression of the Mif2 mutants as only nuclear copies in a Dsn1^S240A,S250A,S264A^ mutant background showed that the Mif2^217-240*9S-A^ mutant was synthetically lethal whereas the Mif2^177-229*9ST-A^ and Mif2^232-240*5S-A^ mutants grew normally (Fig. 5b). The synthetic growth defect of only the phosphorylation-deficient Mif2 mutants, that did not mediate interaction with Ame1/Okp1 *in vitro*, suggests that cooperative kinetochore stabilization through phosphorylation of Dsn1 and the Mif2 region 217-240 is required for cell viability.

The putative Okp1 interface region included 2 predicted helices (Supplementary Fig. 12b and 13b). A deletion mutant of the helix motif Okp1^156-188^, which was previously reported to be essential for binding the Cse4-END (essential-N-terminal-domain)^14^, was lethal but still bound Mif2^*^ *in vitro*, whereas the Okp1^196-229^ helix deletion abrogated Mif2^*^ binding (Fig. 5c and Supplementary Fig. 13c) and inhibited cell growth (Fig. 5d and Supplementary Fig. 15b). Both Okp1 helices form an α-helical hairpin-like structure (Fig. 6)^30,34^ suggesting that the putative phosphorylation of the 9 serines within Mif2^217-240^ establishes a cooperative high-affinity binding environment for the Cse4-NCP by bringing the Mif2^217-240^:Okp1^196-220^, Cse4-END:Okp1^156-188^ and Mif2^285-311^:Cse4^C-term^ contacts into close proximity (Fig. 6 and Supplementary Fig. 11). Moreover, Ame1/Okp1 and Mif2^217-240*9S-A*^ competed for binding to Mtw1/Nnf1 (Fig. 3a)^35^ but formed a nearly stoichiometric complex with *in vitro* phosphorylated wild-type Mif2^*^, suggesting that phosphorylation of the Mif2^217-240^ motif (Fig. 3b) might facilitate the simultaneous stabilization of Mif2^*^ and Ame1 at the same MTW1c (Figs. 5e and 6)^20^.

**Fig. 5:**
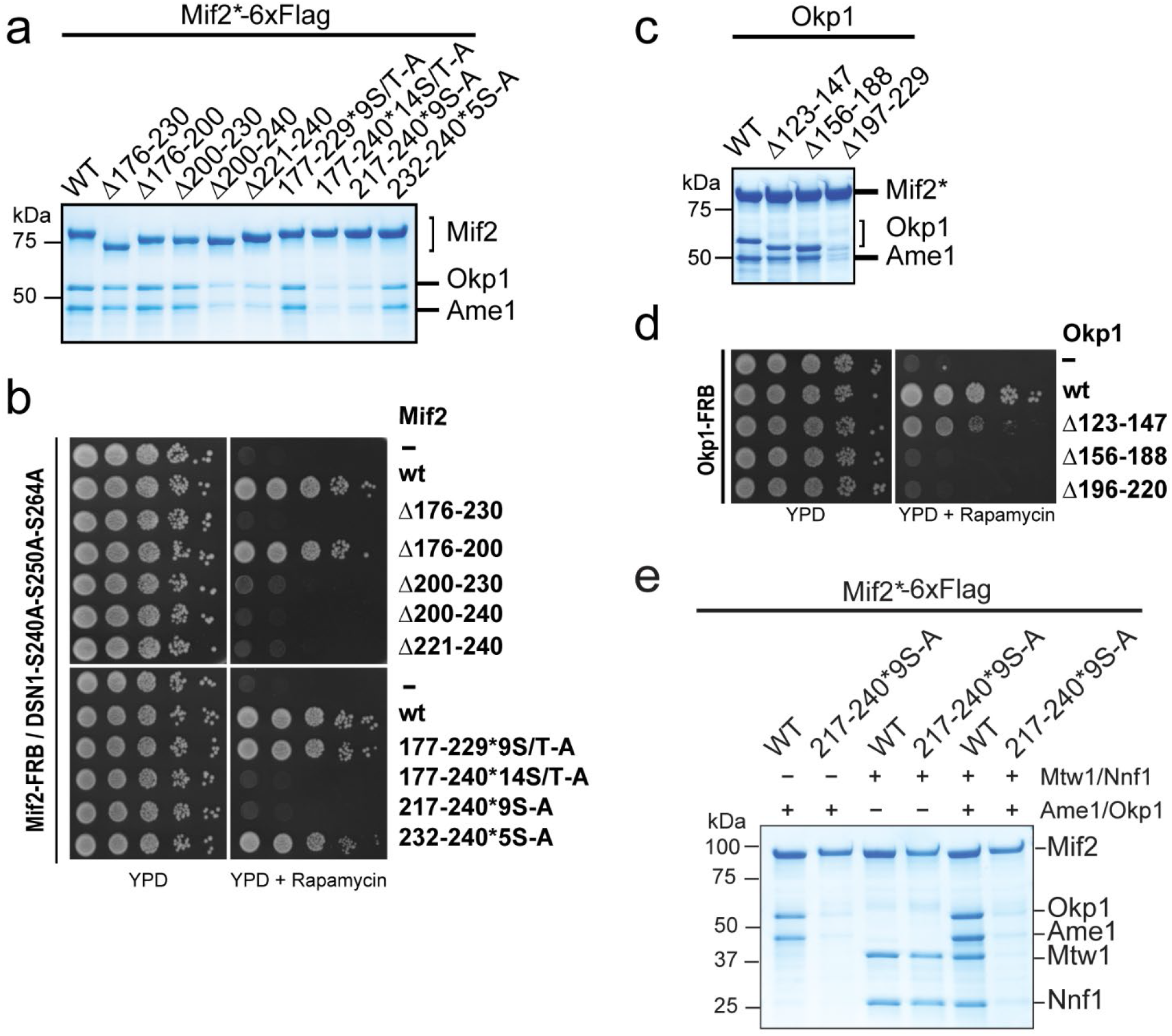
Phosphorylation of Mif2 and Dsn1 mediates a cooperative high-affinity link to the Cse4-NCP and is essential for cell viability. **a**, *In vitro* binding assay to identify the Ame1/Okp1 binding site on Mif2* using the indicated Mif2* deletion and phosphorylation-ablative mutants. **b**, Assay monitoring the rescue of cell growth upon nuclear depletion of Mif2 using the anchor-away method through the ectopic expression of wild-type Mif2 or its indicated deletion or phosphorylation-ablative mutants in a *Mif2-FRB/Dsn1S240A-S250A-S264A* background. **c**, Identification of the Mif2* binding site on Okp1 by assessing the binding of Okp1 deletion mutants *in vitro*. **d**, Assay of the effect of ectopically expressed Okp1 deletion mutants on cell growth in an *Okp1-FRB* anchor-away strain. **e**, *In vitro* assay to determine the effect of 9 putative phosphorylation sites within Mif2^217-240^ on the interaction of Mif2 and Ame1/Okp1 with Mtw1/Nnf1.

**Fig. 6:**
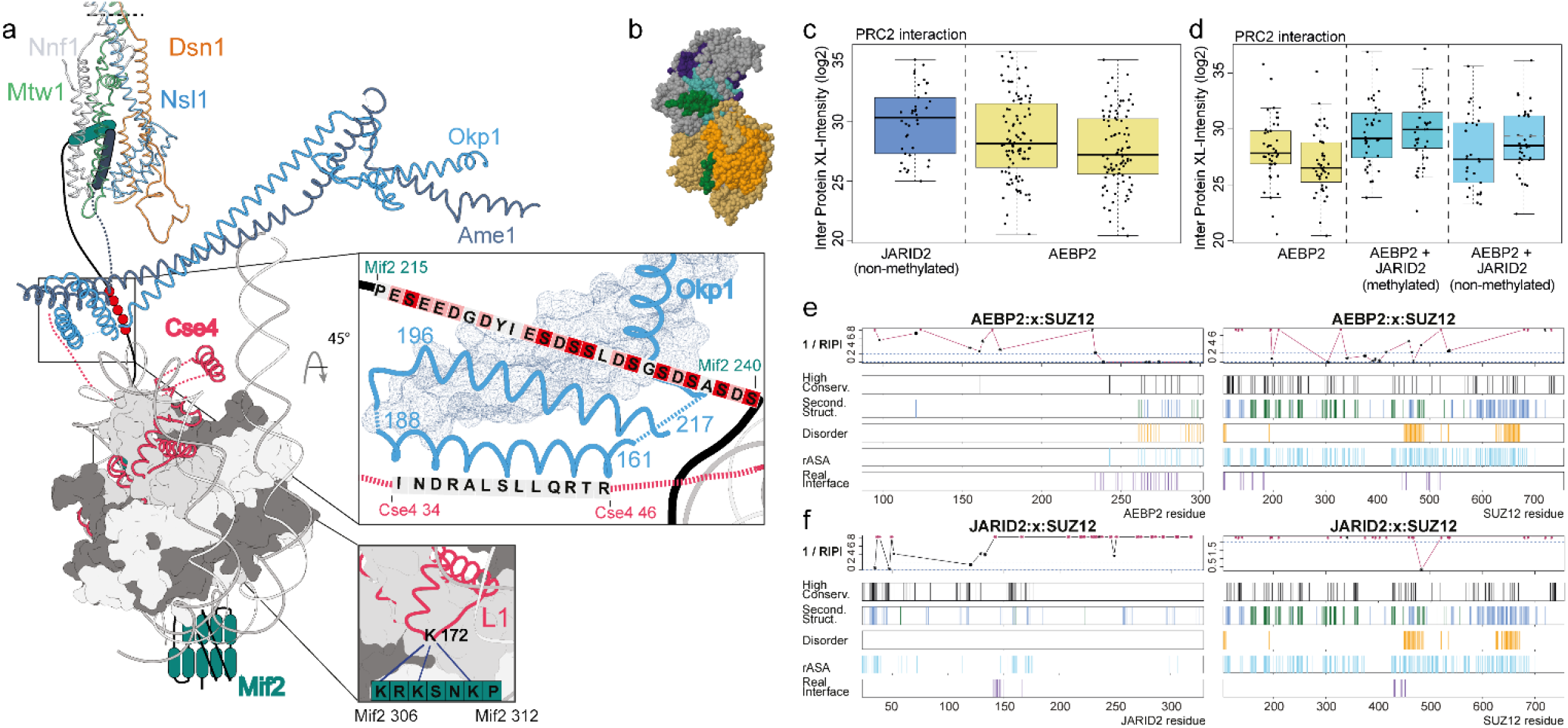
Summary of quantitative XLMS applications to the kinetochore and PRC2 datasets. **a**, Structural model of cooperative kinetochore stabilization on the Cse4 nucleosome through phosphorylation-induced interactions. Model of the MTW1c:Mif2:Ame1/Okp1:Cse4-NCP complex based on cryo electron microscopy and crystal structures (PDB 6NUW, 6QLD, 5T58) depicting the subunit contacts essential for establishing the cooperative binding of Cse4-NCPs by Mif2 and Ame1/Okp1 upon phosphorylation of Dsn1 and Mif2^30,34^. L1 shows Cse4 loop1. Light red and red residues within the Mif2^215-240^ sequence indicate acidic and putatively phosphorylated amino acids, respectively. **b**, Cryo electron microscopy density map of the PRC2 complex with the cofactors JARID2 (dark green) and AEBP2 (cyan) (PDB 6C23) showing the subunits SUZ12 (grey), EED (orange), EZH2 (kaki) and RBAP48 (violet). **c**, Estimation of relative affinities of the cofactors AEBP2 and JARID2 to the PRC2 complex based on crosslink intensities which were extracted and quantified by the TOPP-qXL pipeline. Boxplots with the same colour indicate replicates. **d**, Relative affinity change of AEBP2 for the PRC2 complex in the presence of methylated and non-methylated JARID2. **e, f**, Interface sequence regions are indicated by RIPI blots, calculated from crosslink intensities, for the interactions of SUZ12 with (**e**) AEBP2 and (**f**) JARID2. Inter-protein crosslink lysines are represented as black asterisk. The top 20% conserved residues within the protein sequences are indicated. Secondary structures are shown as alpha helices (blue) and beta strands (green). Real interface residues were obtained from the PDB 6C23.

## Discussion

Our observation that increasing crosslink intensities correlate with shorter crosslink distances lead to the development of a quantitative XLMS approach, which applies inter-protein crosslinks to characterize protein binding interfaces beyond the detection of the protein connectivity. This study demonstrates the capacity of inter-protein crosslink intensities to simultaneously estimate K_D_ values of individual contacts in multi-protein assemblies ranging from 6 to 0.015 µM. Notably, the subset of inter-links proximal to minimal binding interfaces yielded apparent K_D_ values that are in good agreement with values determined by ITC (Figs. 2c and 3d). Moreover, the distance-intensity relation was exploited in the ‘Relative Interface Propensity Index’ to support the prediction of putative interface sequence regions, whose physiological importance was confirmed in cell viability assays.

To demonstrate the applicability of our workflow to datasets, which were not acquired as titration experiments for the purpose of this study, we analyzed the XLMS dataset of the histone H3 methyltransferase Polycomb repressive complex 2 (PRC2) (Fig. 6b)^36^. Based on crosslink intensities we showed that binding of methylated JARID2 increases the relative affinity of the second cofactor AEBP2 to the PRC2 complex (Fig. 6c, d), which is consistent with the observation of a compact active state upon methylation of JARID2 by electron microscopy^36^. In addition, the sequence areas, indicated by the RIPI blot, are in good agreement with the binding interfaces of the PRC2 subunit SUZ12 with the cofactors, JARID2 and AEBP2, which were obtained from electron microscopy density maps (Fig. 6e, f)^36^.

By applying the quantitative XLMS method to analyze the budding yeast kinetochore assembly at centromeric nucleosomes, we identified the interface of the phosphorylation-dependent Mif2:Ame1/Okp1 interaction at the inner kinetochore (Figs. 3a and b). The phosphorylation sites within the Mif2^217-240^ motif established the Mif2:Ame1/Okp1 interaction *in vitro* (Fig. 5a) and were required not only to generate a hub of Cse4 nucleosome binding motifs but might also induce the switch-like stabilization of Mif2 and Ame1 at the outer kinetochore MTW1 complex phosphorylated at the Dsn1 subunit (Figs. 3a, 5e and 6). Together, phosphorylation of the outer kinetochore Dsn1 and the inner kinetochore Mif2 proteins resulted in a ~200-fold increase in Cse4 nucleosome binding affinity *in vitro* (Figs. 3 and 4) and expression of phosphorylation-ablative mutants resulted in synthetic lethality suggesting that the phosphorylation-induced cooperativity is important for kinetochore stabilization *in vivo*. This highlights the capacity of quantitative XLMS to detect the impact of two phosphorylation events on the cooperative stabilization of a macromolecular assembly by a sharp increase in binding affinities.

Although human and budding yeast kinetochores differ in subunit connectivity^8^, the human orthologue of the MTW1 complex, MIS12c, has been implicated in CENP-A stabilization at centromeres^37^. Moreover, we found that the Mif2:Okp1 interface is partially conserved in their human orthologues CENP-C:CENP-Q (Supplementary Fig. 16) and the CENP-C residue T667, which corresponds to Mif2 S226, shows a single nucleotide polymorphism, T667K, in malignant hepatic cancer cells^38^.

We demonstrated that quantitative XLMS facilitated the mechanistic characterization of protein complexes beyond a structural description by estimating protein affinities and their relative changes upon protein modification or ligand interaction. This quantitative XLMS method will significantly contribute to biological modeling at the molecular and cellular level and holds great promise for the development of diagnostic tools for studying the effects of drug interactions on protein complexes and the characterization of epitopes for protein therapeutics.

## Methods

### Protein expression and purification of Spc24/25, MTW1c, Cnn1^1-270^, Ame1/Okp1, Clb2 and Mps1 from *E. coli*

For the expression of the budding yeast Spc24/25 complex in *E. coli*, the respective genes were amplified from genomic DNA and cloned into the pETDuet-1 vector (Novagen). Expression and purification of the Spc24/25 complex were performed as described previously ^15^. In brief, pETDuet1-Spc24-6xHis/Spc25 was transformed into *E. coli* strain BL21 DE3 (EMD Millipore). Bacteria were grown in selective LB-medium to an OD_600_ of 0.6 at 37 °C and protein expression was induced with 0.2 mM IPTG for 18 h at 18 °C. Cells were lysed in lysis buffer (30 mM HEPES, pH 7.5, 300 mM NaCl, 5% glycerol, 30 mM imidazole, Complete EDTA-free protease inhibitor [Roche]) and the cleared lysate was incubated with Ni-NTA agarose beads (Qiagen). The protein complex was eluted with buffer containing 30 mM HEPES pH 7.5, 150 mM NaCl, 0.01% NP40, 2% glycerol and 250 mM imidazole and further purified on a Superdex 200 HiLoad 16/600 column (GE Healthcare) in the gel filtration buffer (30 mM HEPES pH 7.5, 150 mM KCl and 5% glycerol).

The constructs for budding yeast Mtw1/Nnf1 (pETDuet-Mtw1-Nnf1-6xHis) and Dsn1/Nsl1 (pST-39-Mtw1-Nsl1-6xHis-Dsn1) were kindly provided by S. Westermann ^15^. For the phospho-mimetic version of MTW1c (Mtw1/Nnf1/Dsn1^S240DS250D^/Nsl1), the serine residues S240 and S250 in Dsn1 were mutated to aspartic acid using the Q5 site-directed mutagenesis kit (New England Biolabs) as described previously ^19,20^. The plasmid containing Mtw1/Nnf1 was transformed into *E. coli* Rosetta (DE3) strain (EMD Millipore), whereas Dsn1/Nsl1 was transformed into BL21 DE3 (EMD Millipore). Transformed bacteria were grown in selective LB medium at 37 °C to OD_600_ 0.6-0.8 and protein expression was induced with 0.2 mM IPTG (Mtw1/Nnf1 expression) or 0.5 mM IPTG (Dsn1/Nsl1) at 18 °C for 18 h. Cells were lysed in lysis buffer (50 mM HEPES, pH 7.5, 400 mM NaCl, 5% glycerol, 20 mM imidazole, 1 Mm DTT, Complete EDTA-free protease inhibitor [Roche]) and the cleared lysate was incubated with Ni-NTA agarose beads (Qiagen). After several washing steps in wash buffer (50 mM HEPES, pH 7.5, 600 mM NaCl, 5% glycerol, 20 mM imidazole) the protein complex was recovered in elution buffer (50 mM HEPES pH 7.5, 150 mM NaCl, 5% glycerol, 300 mM imidazole). To reconstitute the MTW1c, fractions containing pure protein sub-complexes were subjected to size-exclusion chromatography (Superose 6 increase 10/300, GE Healthcare) in 25 mM HEPES pH 7.5, 150 mM KCl, 5% glycerol and fractions containing reconstituted MTW1c were collected, flash-frozen in liquid nitrogen and stored at −80 °C.

The construct encoding Ame1-6xHis/Okp1 (pST39-Okp1-Ame1-6xHis) was kindly provided by S. Westermann ^15^. Protein expression and purification in *E. coli* was essentially performed as described ^15^ with the modification that 25 mM HEPES buffer was used as buffer component in all purification steps and the final gel filtration was performed on a Superdex 200 HiLoad 16/600 column (GE Healthcare) in 25 mM HEPES pH 7.5, 150 mM KCl, 5% glycerol.

For Mps1 expression and purification, the Mps1 coding sequence was cloned into pETDuet-1 with an N-terminal 6xHis-tag. Protein expression and purification was performed as described for the MTW1c and the Ni-NTA eluate was desalted using a PD10 column (GE Healthcare) in desalting buffer (50 mM HEPES pH 7.5, 150 mM NaCl, 10% glycerol, 0.5 mM DTT).

The construct for budding yeast Cnn1^1-270^ (pETDuet-6xHis-Cnn1^1-270^) was kindly provided by S. Westermann ^28^ and purified as described. After elution, the protein was further purified on a Superdex 200 HiLoad 16/600 column (GE Healthcare) in gel filtration buffer (25 mM HEPES pH 7.5, 150 mM KCl and 5% glycerol).

### CDC28^CDK1^ complex purification

Reconstitution of the CDC28 complex, consisting of Clb2, Cdc28 and Cks1, could not be performed by the single expression of all partners from a single baculovirus in insect cells, as Clb2 was degraded. To reconstitute the three subunit CDC28c, 1xStrep-tagged Clb2 was expressed and purified from *E. coli*, immobilized on Strep-Tactin beads (Qiagen) and incubated with cell lysate of baculovirus infected High Five™ cells expressing Cdc28 and Cks1, to assemble the three subunit CDC28c.

Full-length Clb2 was PCR amplified from budding yeast genomic DNA and cloned into pET-28 with an N-terminal 1xStrep-tag. pET28-1xStrep-Clb2 transformed *E. coli* Rosetta (DE3) (EMD Millipore) cells were grown in selective LB medium to OD_600_ 0.6-0.8 and expression of Clb2 was induced by 0.4 mM IPTG at 18 °C for 18 h. Cells were resuspended in lysis buffer containing 50 mM HEPES pH 7.5, 150 mM KCl, 5% glycerol, 0.01% Tween, 1.5 mM MgCl_2_, 1 mM DTT and complete EDTA-free protease inhibitor (Roche) and lysed by sonication. The cleared lysate was incubated with Strep-Tactin Superflow resin (Qiagen) for 1 h at 4 °C. Immobilized Clb2 was washed with wash buffer (50 mM HEPES pH 7.5, 150 mM KCl, 5% glycerol, 1 mM DTT) and incubated with the cleared insect cell lysates containing recombinant Cdc28 and Cks1 for one hour at 4 °C. Beads were washed and the reconstituted CDC28 complex was recovered in elution buffer (50 mM HEPES pH 7.5, 300 mM KCl, 5% glycerol, 1 mM DTT, 10 mM biotin). The eluate was dialyzed in 50 mM HEPES pH 7.5, 150 mM KCl, 10% glycerol) and flash-frozen aliquots were stored at −80 °C.

### Protein expression and purification from insect cells

Open reading frames encoding the respective subunits were amplified from yeast genomic DNA and cloned into the pBIG1/2 vectors for insect cell expression according to the biGBac protocol ^39^. Generation of recombinant viruses expressing single or multiple subunits was performed according to the MultiBac system ^40^.

Mif2-6xHis-6xFlag wild-type and mutant proteins were expressed in High Five™ cells for three days at 27 °C. Cells were lysed in lysis buffer (30 mM HEPES pH 7.5, 400 mM NaCl, 20 mM imidazole, 5% glycerol, 125 U/ml benzonase (Merck), 1 mM MgCl_2_ and complete protease inhibitor cocktail [Roche]) using a dounce homogenizer. The cleared lysate was incubated with Ni-NTA resin (Qiagen) washed with lysis buffer (without protease inhibitor) and eluted in 30 mM HEPES pH 7.5, 150 mM NaCl, 5% glycerol and 250 mM imidazole.

6xHis-Cdc5^Plk1^ was expressed and purified from insect cells as described for Mif2 with the following modifications. Cells were lysed in lysis buffer (50 mM HEPES pH 7.5, 150 mM NaCl, 5% glycerol, 125 U/ml benzonase (Merck), 1 mM MgCl_2_ and complete protease inhibitor cocktail [Roche]) using a dounce homogenizer. The cleared lysate was incubated with Ni-NTA resin (Qiagen), washed with 50 mM HEPES pH 7.5, 300 mM NaCl, 20 mM imidazole, 5% glycerol and eluted in 50 mM HEPES pH 7.5, 150 mM NaCl, 5% glycerol and 250 mM imidazole. Peak fractions were combined and the buffer was exchanged using a PD10 column (GE Healthcare) in desalting buffer (50 mM HEPES pH 7.5, 120 mM NaCl, 3% glycerol).

Sli15ΔN228-2xStrep/Ipl1 complex was purified from insect cells as described previously ^14^. Insect cell lysates containing expressed untagged Cdc28 and Cks1 were prepared as described above in lysis buffer (50 mM HEPES, pH 7.5, 150 mM KCl, 5 % glycerol, 0.01% Tween and complete EDTA-free protease inhibitors [Roche]) and the cleared lysates were used to assemble the trimeric CDC28 complex with 1xStrep-Clb2 purified from *E. coli*.

For *in vitro* binding and quantitative crosslinking experiments Cdc5^Plk1^ phosphorylated Mif2 was generated according to the following procedure. 1 mg 6xHis-tag purified Mif2-6xHis-6xFlag was immobilized on anti-FlagM2 agarose beads (Merck) for 1 h, at 4 °C. Unbound protein was removed by washing 2x with wash buffer (30 mM HEPES pH 7.5, 150 mM NaCl, 5% glycerol). Subsequently, Mif2 was treated for 2 h at 30 °C with lambda-phosphatase (*New England Biolabs)* according to the manufacturer’s instruction. The dephosphorylation reaction was stopped by washing 1x in wash buffer supplemented with Halt™ Phosphatase Inhibitor Cocktail (Thermo Fisher) and 2x without phosphatase inhibitors. Mif2 was re-phosphorylated by adding 50 µg Cdc5^Plk1^ in the presence of 2.5 mM MgCl_2_ and 1 mM ATP at 30 °C. The kinase reaction was stopped by washing 2x in wash buffer and Mif2 was recovered in elution buffer (30 mM HEPES pH 7.5, 150 mM NaCl, 5% glycerol, 1 mg/ml 3xFLAG-peptide). For quantitative crosslinking experiments the eluate was further purified on a Superdex 200 HiLoad 16/60 column (GE Healthcare) in gelfiltration buffer (30 mM HEPES pH 7.5, 150 mM KCl and 5% glycerol).

### *In vitro* binding assay of Mif2 wild-type and mutant proteins to Ame1/Okp1

To analyze the interaction of Ame1-6xHis/Okp1 with Mif2-6xHis-6xFlag wild-type and mutant proteins *in vitro*, 10 µM Cdc5^Plk1^ re-phosphorylated Mif2 protein (M3) was immobilized on anti-FlagM2 beads (Merck) and incubated with 25 µM Ame1/Okp1 complex in binding buffer (50 mM HEPES pH 7.5, 150 mM NaCl, 3% glycerol, 0.01% Tween 20) for 1 h at 4 °C and 1200 rpm in a thermomixer (Eppendorf). Unbound protein was removed by washing 2x with high salt buffer (50 mM HEPES pH 7.5, 300 mM NaCl, 3% glycerol, 0.01% Tween 20) and 1x with binding buffer. Bound protein was eluted in binding buffer containing 1 mg/ml 3xFLAG peptide (Ontores).

To test the binding of Mif2 and Ame1/Okp1 to Mtw1/Nnf1, 10 µM Mif2-6xHis-6xFlag or Mif2 S217-240A-6xHis-6xFlag was incubated with 20 µM Mtw1-Nnf1-6xHis and immobilized on anti-FlagM2 beads (Merck) for 1 h at 4 °C and 1200 rpm. The beads were washed 1x with high salt buffer and 1x with binding buffer. The complex was subsequently incubated with 10 µM Ame1/Okp1 complex in binding buffer for 1 h at 4 °C and 1200 rpm. Unbound Ame1/Okp1 was removed by washing 2x with high salt buffer and 1x with binding buffer. Proteins were eluted in a buffer containing 50 mM HEPES pH 7.5, 150 mM NaCl, 5% glycerol and 1 mg/ml 3xFLAG peptide (Ontores). The input and bound fractions were separated by SDS-PAGE and proteins were visualized by Coomassie brilliant blue staining.

To analyze the binding of untreated, dephosphorylated or re-phosphorylated Mif2-6xHis-6xFlag wild-type to Ame1-6xHis/Okp1 *in vitro*, 10 µM Mif2 protein per condition was immobilized on anti-FlagM2 agarose beads (Merck) for 1 h at 4 °C and 1200 rpm in a thermomixer. The beads were washed 3x with wash buffer (50 mM HEPES pH 7.5, 150 mM NaCl, 3% glycerol, 0.01% Tween 20) and an aliquot of the untreated sample was removed. Anti-Flag immobilized Mif2-6xHis-6xFlag was then treated with lambda-phosphatase (*New England Biolabs)* according to the manufacturer’s instruction and incubated for 2 h at 30 °C and 1200 rpm in a thermomixer. The dephosphorylation reaction was stopped by washing 1x in wash buffer supplemented with Halt™ Phosphatase Inhibitor Cocktail (ThermoFisher) and 2x without phosphatase inhibitors. An aliquot of the lambda-phosphatase treated sample was removed and the rest was aliquoted and used in *in vitro* kinase assays with CDC28c, Cdc5, Sli15/Ipl1, Mps1 or combinations thereof in the presence of 2.5 mM MgCl_2_ and 1 mM ATP for 30 min at 30 °C and 1200 rpm. The kinase reaction was stopped by washing 1x with high salt buffer and 2x with wash buffer. The binding of the untreated, dephosphorylated and re-phosphorylated Mif2-6xHis-6xFlag samples to Ame1-6His/Okp1 was analyzed as described. Quantification of the ratios of bound protein to the bait was performed by using ImageJ ^41^ from three independent experimental set-ups.

### *In vitro* reconstitution of Cse4- and H3-containing nucleosome core particles (NCPs)

Octameric Cse4 and H3 containing nucleosomes were *in vitro* reconstituted from budding yeast histones which were recombinantly expressed in *E. coli* and assembled on the 147 bp ‘Widom601’ nucleosome positioning sequence according to a modified protocol ^42,43^.

### Protein complex titration, chemical crosslinking and mass spectrometry

The purified proteins and protein complexes were titrated applying a series of molar ratios and incubated for 45 min at room temperature to allow complex formation. For example, the titration of the Cnn1^1-270^-Spc24/25 complex was performed by incubating Cnn1^1-270^ with the Spc24/25 dimer at molar ratios of 0.05, 0.15, 0.25, 0.55, 0.60, 0.65, 0.75, 0.80, 0.85, 0.90, 0.95, 1.0, 1.25, 1.5 and 2.0 in a final volume of 95 µl at 25 °C. Subsequently, protein complexes were crosslinked by the addition of an equimolar mixture of isotopically light (hydrogen) and heavy (deuterium) labelled bis(sulfosuccinimidyl) 2,2,4,4-glutarate (BS2G-d_0_/d_6_) (Creative Molecules) at a final concentration of 0.5-0.75 mM at 30 °C for 2 min. The crosslinking reaction was quenched by adding ammonium bicarbonate to a final concentration of 100 mM for 20 min at 30 °C. Proteins were diluted by adding 2 volumes of 8 M urea, reduced by 5 mM TCEP (Thermo Fisher) at 35 °C for 15 min and alkylated by incubating with 10 mM iodoacetamide (Sigma-Aldrich) at room temperature for 30 min in the dark. Proteins were digested with Lys-C (1:50 (w/w), Wako Pure Chemical Industries) for 2 h at 35 °C and 1300 rpm, diluted to 1 M urea with 50 mM ammonium bicarbonate and digested with trypsin (1:50 (w/w), Promega) overnight at 35 °C and 1300 rpm. Peptides were acidified by adding trifluoroacetic acid to a final concentration of 1% and purified by reversed phase chromatography using C18 cartridges (Sep-Pak, Waters). Crosslinked peptides were enriched by size exclusion chromatography on a Superdex Peptide PC 3.2/30 column (GE Healthcare) using water/acetonitrile/TFA (77.4/22.5/0.1, v/v/v) as mobile phase at a flow rate of 50 μl/min. Fractions containing crosslinked peptides were analyzed by liquid chromatography coupled to tandem mass spectrometry (LC-MS/MS) using an EASY-nLC 1200 and an LTQ-Orbitrap Elite mass spectrometer (Thermo Fisher). Peptides were injected onto a 15 cm x 0.075 mm i.d. Acclaim™ PepMap™ C18 column (2 µm particle size, 100 Å pore size) and separated at a flow rate of 300 nl/min using the following gradient: 0-5 min 3% B and 5-65 min 3-35% B (acetonitrile/water/formic acid, 98:2:0.1). The mass spectrometer was operated in data-dependent mode, selecting up to 10 precursors from a MS1 scan (resolution 60,000) in the range of m/z 350–1800 for collision-induced dissociation excluding singly and doubly charged precursor ions and precursors of unknown charge states. Dynamic exclusion was activated with a repeat count of 1, exclusion duration of 30 s, list size of 300, and a mass window of ±50 ppm. Fragment ions were detected at low resolution in the linear ion trap.

### Identification of peptide crosslink spectra

Raw spectra were converted to mzXML format using MSConvert ^44^ and crosslink spectra were searched and identified using xQuest/XProphet ^24^. Peptide spectrum matches were performed against a database including the subunits of the respective complex (Spc24, Spc25, Cnn1 or Mif2, Ame1/Okp1, Cse4-NCP, Mtw1/Nnf1/Dsn1/Nsl1) and 22 *E. coli* decoy protein sequences. A maximum of two trypsin missed cleavages and peptide lengths between 4 and 45 amino acids were allowed. Carbamidomethyl-Cys was set as a fixed modification and a mass shift of 96.0211296 for intra-/inter-protein crosslink candidates with an additional shift of 6.03705 to account for crosslinks with the heavy version of BS2. A precursor mass tolerance of ±10 ppm was used and a tolerance of 0.2 and 0.3 Da for linear and crosslinked fragment ions, respectively. The search was performed in the ‘ion-tag’ mode. Identifications were filtered by applying a maximum FDR of 5%, precursor errors of ±5.0 ppm, a maximum delta score of 0.9 and a minimum of 3 fragment ion matches per peptide. The final identification tables were downloaded as xtract.csv files from the xQuest/xProphet visualization tool.

### Quantification of peptide-peptide crosslinks and site-site crosslinks using the TOPP-qXL pipeline

Quantification was performed with an in-house developed workflow based on the OpenMS software version 2.0 ^25^. All scripts as well as the xtract.csv files to run the python script ‘toppXLquant.py’ (C:/Users/…/Scripts/TOPPqXL/bin/) are provided in the ‘Scripts.zip’ folder. The pipeline starts with the conversion of the identification tables in the xtract.csv files to idXML format using our script ‘xtractToIdXML.py’. The files were saved and the workflow ‘basic_xlquant.toppas’ (C:/Users/…/Scripts/TOPP-qXL/workflows) was opened in the OpenMS framework ^25^. The ‘*.idXML’ and ‘*.mzXML’ files are uploaded as input files of the workflow. During execution of the workflow, raw files in the mzXML format were converted to mzML using the FileConverter function with default parameters except for the filtering of MS2 scans and MS1 peaks with intensities <100.0. Peak features in the mzML files and their respective profile chromatograms were extracted with the FeatureFinderAlgorithmPicked function from OpenMS. Parameters fed to this tool are found in the file ‘ffcentroided_params.ini’. Detected features were annotated with their putative peptide identifications in the idXML files using the IDMapper function with an m/z tolerance of ±7 ppm and RT tolerance of ±10 s. Retention times between runs were aligned using the MapAlignerIdentification function with default parameters. Finally, consensus tables were generated using the FeatureLinkerUnlabeled function with default parameters and converted to .csv format with the TextExporter function. The intensities of the unique peptide-peptide crosslink ions were summarized to site-site crosslink intensities using the in-house script ‘csvToToppXLqTSV.py’ (provided in: C:/Users/…/ Scripts/TOPPqXL/bin).

### Estimation of the apparent equilibrium dissociation constant (K_D_) based on crosslink intensities

Site-site crosslink intensities were loaded and analyzed in the statistical environment R (https://www.r-project.org). Technical replicates were averaged with non-assigned values being ignored at this step. The intensities of peptides seen in >1 SEC fraction were summed up and peptide-peptide crosslinks were summarized to site-site crosslinks by addition of their intensities. The intensities of the subunit whose concentration was constant in all titrations were applied to normalize the intensities between runs. Finally, a linear model was fitted between the initial concentrations of the varying subunit and the median intensity of its intra-protein crosslinks. This linear relation was used to estimate the concentration of the formed complex from the median intensity of the inter-protein crosslinks. Subsequently, the K_D_ was calculated as:

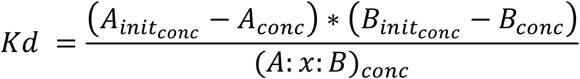

where *A* represents the subunit whose concentration varies, *B* the subunit whose concentration remains constant and *A*:x:*B* the complex. The initial concentrations of *A* and *B* were recalculated based on the linear relation of concentration and intensity. For each titration step a K_D_ value was calculated and the mean and standard deviation of these values were reported. We also applied the *Scatchard plot* ^29^ to estimate the K_D_ by plotting the linear relation of ‘fraction of B bound over concentration of free A’ (y-axis) versus ‘fraction of B bound’ (x-axis). This approach indicates the K_D_ as the negative inverse of the slope as well as the inverse of the intersection coefficient (Fig. 2b).

To calculate the apparent K_D_ values based on the steady state equilibrium equation the R script was run according to the following procedure. The scripts (C:/Users/…/Scripts/R-Script) were opened in the R environment. To analyze the Cnn1:x:Spc24/25 titration the ‘CnnSPC_Kd_Est.R’ script and for the analysis of the Mif2:Ame1/Okp1:MTW1c:x:Cse4-NCP titration the ‘MTW1cMifAO_CSE4-NCP_Kd_Est.R’ script were applied. The location of the input files was defined in the working directory in setwd(“C:/Users/…/”). The input file name was defined in ‘fname’ (e.g.: fname = “1.1-MIFNUC_F restraints.tsv”). Subsequently, the default settings of the calculation parameters, as described above, can be altered by following the instructions in the code. Executing the script shows the results table (‘kdtable2’) which indicates the K_D_ values of each titration step and the mean (KD) and standard deviation (SD). At this step outliers that exceed the double SD are excluded and the mean K_D_ (KD2) and standard deviation (SD2) are recalculated. In addition, several exploratory plots are generated. (1) Crosslink intensities per protein:x:protein pair (median) before normalization (Supplementary Figs. 4a, 5, 8a, 8c, 8e, 8g and 10a). (2) Correlation of crosslink intensities within protein:x:protein pairs. (3) Crosslink intensities per protein:x:protein pair (median) after normalization (Supplementary Figs. 4b, 8b, 8d, 8f, 8h and 10b). (4) Correlation of crosslink intensities between experiments and between crosslinks. (5) Linear regression between crosslink intensity and protein concentration. The linear regression model is used to estimate the apparent K_D_ values. The statistical analysis of the apparent K_D_ values for each interaction is summarized in ‘kdtable2’.

### Determination of the Relative Interface Propensity Index (RIPI)

Peptide-peptide crosslink intensities were summarized to site-site intensities, by summing up all restraint intensities involving the specific lysine residue. This total sum includes mono-links, loop-links, intra- and inter-protein crosslinks. Next, the site-site intensity of the inter-protein crosslinks from a specific dimer interaction was divided by the total sum. The resulting value was called the Relative Interface Propensity Index (RIPI) of a crosslinked residue. Lysine sites, which were not identified in inter-protein crosslinks, were assigned a RIPI value equal to the minimum RIPI in the set, in order to avoid infinite values for the plotted inversed RIPIs.

Sequence conservation in the RIPI plots was computed by using PSIBlast against the UNIREF90 database. Only residue positions with conservation above the 80% quantile within the protein sequence were plotted.

Secondary structure and rASA (relative accessible surface area) were predicted using the SPIDER2 software ^45^ against the UNIREF90 database. The fasta protein sequences and the PSSMs (Position-Specific Scoring Matrix) obtained by PSIBlast were used as input for the SPIDER2 software. Residues were considered to have low accessibility if their rASA was below 40%. Residues were considered to have low disorder if their IUPred index was below 0.25 in a scale of 0 to 1.

Real interface residues were extracted from PDB models if applicable. Real binding interfaces were identified by a residue-residue distance between the interacting proteins of below 4.5 Å. The distances were measured from any heavy atom in one residue to any heavy atom in the other residue.

### Yeast strains and methods

All yeast strains used in this study were created in the S288c background and are listed in Supplementary Table 8. The generation of yeast strains and yeast methods were performed by standard procedures. The anchor-away analysis was performed as described previously ^46^.

For anchor-away rescue experiments, the Mif2 promoter (1 kb) and coding sequence were PCR amplified from yeast genomic DNA and cloned with a 6xHis-7xFlag tag PCR fragment into vector pRS313 via the Gibson assembly reaction ^47^. The deletion mutants were generated using the Q5 site-directed mutagenesis kit (New England Biolabs) and phospho-ablative mutants were constructed by Gibson assembly of the corresponding mutant gene fragments (IDT). The rescue constructs were transformed into a Mif2 anchor-away strain (*Mif2-FRB*) or a *Mif2-FRB/dsn1*^*S240AS250AS264A*^ mutant strain (Supplementary Table 8) and cell growth was tested in 1:10 serial dilutions on YPD plates in the absence or presence of rapamycin (1 mg/ml) at 30 °C for 3 days.

### Western blot analysis

The levels of proteins ectopically expressed in yeast were probed by western blot analysis as described previously ^14^. For western blot analysis an equivalent of 10 OD_600_ of cells logarithmically grown in selective liquid culture was collected by centrifugation at 3140 x g for 5 min at room temperature and the pellet was washed once with aqua dest. For protein extraction, the pellet was resuspended in 1 ml ice-cold 10% trichloroacetic acid and incubated on ice for 1 h. Samples were pelleted at 4°C and 20000x g for 10 min and washed twice with ice-cold 95% ethanol. Pellets were air-dried and resuspended in 100 µl 1x SDS-PAGE sample buffer containing 75 mM Tris (pH 8.8). Samples were boiled (10 min, 95°C) and centrifuged at 10800 x g for 3 min at room temperature and supernatants were separated on 10% SDS-PAGE gels. Immunoblotting was performed with Anti-FLAG M2 (Sigma-Aldrich) or Anti-PGK1 (ThermoFisher) antibodies and visualized by HRP-conjugated anti-mouse secondary antibodies (Santa Cruz).

### Amino acid sequence alignment

Multiple sequence alignments of *S. cerevisiae* Mif2 and Okp1 amino acid sequences with their respective mammalian orthologues CENP-C or CENP-Q were performed with Clustal Omega^48^ (https://www.ebi.ac.uk/Tools/msa/clustalo/).

## Acknowledgements

We thank Ruedi Aebersold, Stefan Westermann and Alexander Leitner for comments on the manuscript. GH and CK were funded by the Graduate School GRK 1721 and MP was supported by the Graduate School Quantitative Biosciences Munich of the German Research Foundation (DFG). FH was supported by the European Research Council (ERC-StG no. 638218), the Human Frontier Science Program (RGP0008/2015), by the Bavarian Research Center of Molecular Biosystems and by an LMU excellent junior grant.

## Data and materials availability

The mass spectrometry raw data was uploaded to the PRIDE Archive. The access information for reviewers is Project Name: Quantitative Crosslinking and Mass Spectrometry Detects Phosphorylation-Induced Kinetochore Stabilization, Project accession: PXD020094, Username, Password: JFeuElbD.

